# Biological Perspective of Balur Therapy: A Network Pharmacology Study

**DOI:** 10.1101/2020.11.05.366039

**Authors:** Sherry Aristyani, Saraswati Subagjo, Tintrim Rahayu, Sutiman Bambang Sumitro

## Abstract

Balur is an integrative topical medication from Indonesia using herbal medicines: *Moringa oleifera*, *Nicotiana tabacum*, and *Coffea arabica*. Balur can be applied to treat various diseases including chronic diseases and autism because it performs to scavenge free radicals and release electron of heavy metals. However, the complexity of the balur mechanism as medical therapy needs a comprehensive understanding. Not only from a modern physics perspective but also from a biological perspective to explore the effect of active compounds on the human body. In this study, we proposed the computational study to understand balur therapy from a biological perspective though the molecular mechanism. Active compounds of 3 herbal medicines Balur were collected from Dr. Duke’s Phytochemical and Ethnobotanical Databases. Proteins target related to active compounds were obtained from SwissTargetPrediction and PharmMapper Server. Gene Ontology (GO) was conducted to verify the potential mechanism. Moreover, network analysis was conducted with Cytoscape. We found that the active compounds were contributed to the therapeutic effectiveness through a molecular mechanism. This study demonstrated the multi-compounds and multi-target of balur’s herbal medicines to treat disease.

## Introduction

Excessive free radicals in the human body lead to chronic and degenerative diseases (Pham-huy et al., 2008). Free radicals result in protein oxidation by hydrogen abstraction, electron transfer, addition, fragmentation, and rearrangement reaction with amino acid, peptide, and protein, thus causing cellular damage and protein degradation (Davies, 2016). Free radicals are also induced DNA mutation by DNA oxidative damage through abstractions and addition reactions of free radicals to the nitrogenous bases of DNA and the sugar moiety (Dizdaroglu et al., 2002; Dizdaroglu & Jaruga, 2011). Balur is one of the topical complementary and alternative therapy from Indonesia which can release free radicals from the human body by disjoining free radicals bound within aromatic amino acid and DNA by chelation process.

In the process of balur, the patients have lied on a grounded copper table to cascade negative free electrons into the earth generated during the balur process. First, the whole body of the patient is smeared with acetosal liquid to open the skin pores, then smeared with *Moringa oleifera’*s leaves fermentation, condensed, and covered with aluminum foil to induce body warm and release free radicals bound with an active compound from herbal medicines through skin. Next, a mixture of hydroxyurea and mannitol is smeared on the body to collect the free radicals, then warm hydroxy urea liquid is tapped to pull out free radicals through the skin, and the body is covered again with aluminum foil. In the last step, the body is smeared with coffee. For all smearing, the process is applied counterclockwise.

Balur is not only for treating multiple chronic and degradative diseases but also for improving the quality of life. Therefore, comprehensive knowledge is needed to understand the complex mechanisms of balur in the human body. This study tried to know balur in computational biology way by exploring the target proteins of *Moringa oleifera*, *Nicotiana tabacum*, and *Coffea arabica* used in balur.

## Method

The flow chart of the procedure is shown in Figure 1. The active compounds of *Moringa oleifera*, *Nicotiana tabacum*, and *Coffea arabica* were collected from Dr. Duke’s Phytochemical and Ethnobotanical Databases (https://phytochem.nal.usda.gov/). Active compounds were selected according to part of the plant used in the balur process: leaf for *Moringa oleifera* and *Nicotiana tabacum*; seed for *Coffea arabica.* SwissTargetPrediction (http://www.swisstargetprediction.ch) with probability ≥ 0.6 and PharmMapper (http://www.lilab-ecust.cn/pharmmapper/) with fit score ≥ 0.6 were used to collecting putative target proteins of active compounds. For exploring the mechanism of target proteins and predictive disease, Gene Ontology (GO) with PANTHER (http://pantherdb.org) and KEGG disease database (https://www.genome.jp/kegg/disease/) was used. Cytoscape 3.8.1 was used to visualize the network of active compounds-target proteins and target proteins-predictive disease.

**Figure. 1.**
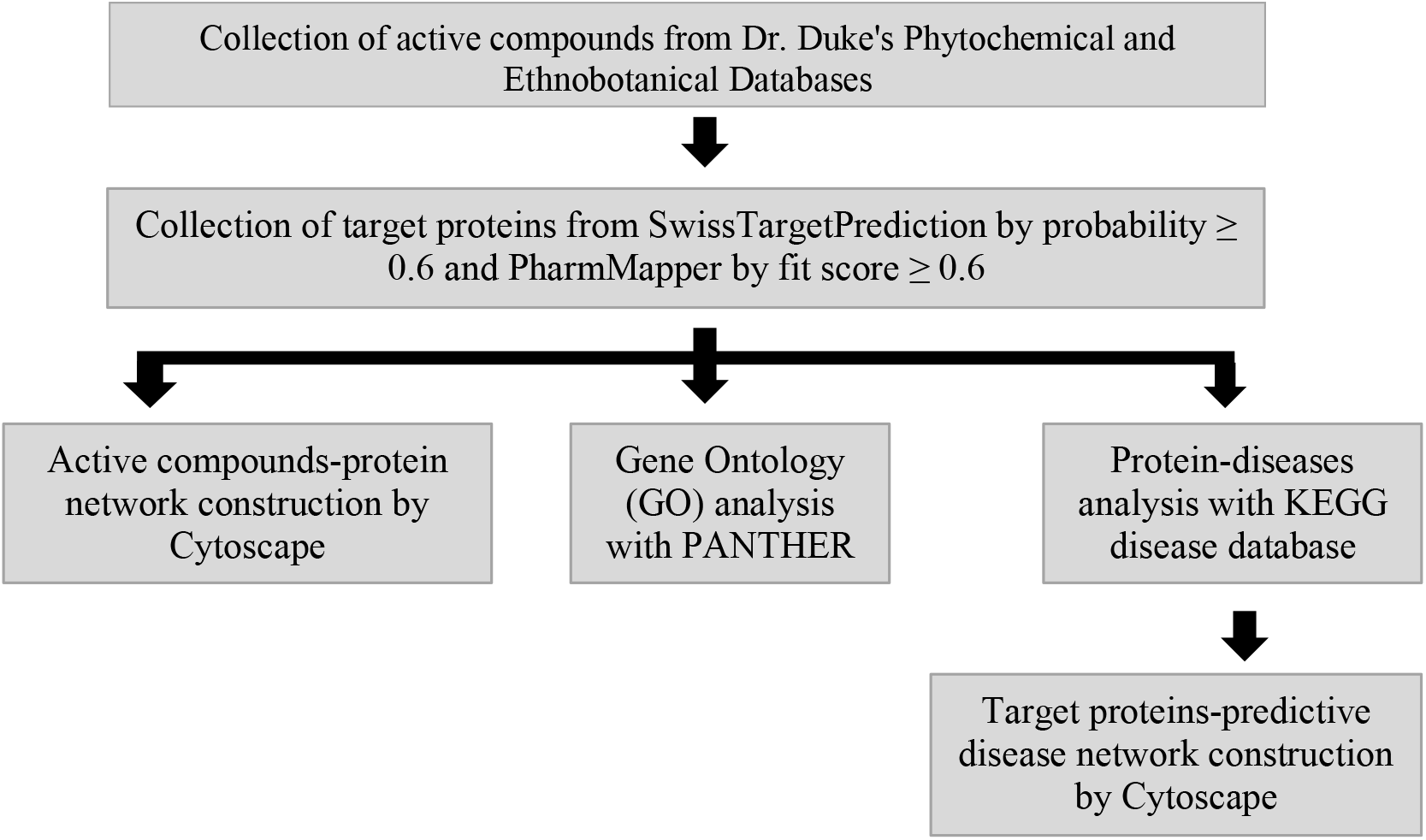
Research procedure’s flow chart

## Results and Discussion

Active compounds obtained from Dr. Duke’s Phytochemical and Ethnobotanical Database were total 383 consisting: 25 of *Moringa oleifera*’s leaf, 228 of *Nicotiana tabacum*’s leaf, and 130 of *Coffea arabica*’s seed. According to the target proteins searching result, it was found 67 compounds were 252 targeted human proteins, which phytate, quercetin, xylan, kaempferol, and quercitrin were the top five active compounds binding numerous (Table 1). Phytate or phytic acid, a compound found in grains and seeds of plants, is an antioxidant agent which uses a chelating iron mechanism to inhibit hydroxyl radicals and suppress linoleic acid oxidation and the formation of 4-hydroxyalkenals, a lipid peroxidation product (Rimbach & Pallauf, 1998; Zajdel et al., 2013). Quercetin, kaempferol, and quercitrin belong to the flavonoid group, a large group plant’s secondary metabolite. Flavonoid has an antioxidant role as inhibiting enzyme or chelating elements to reduce free radical formation, scavenging free radicals, and protecting antioxidant defenses (Kumar & Pandey, 2013). Xylan is abundant biopolymer hemicellulose, and it has been reported that xylan has 70% iron-chelating activity as an antioxidant mechanism (Melo-Silveira RF et al., 2012).

**Table 1.**
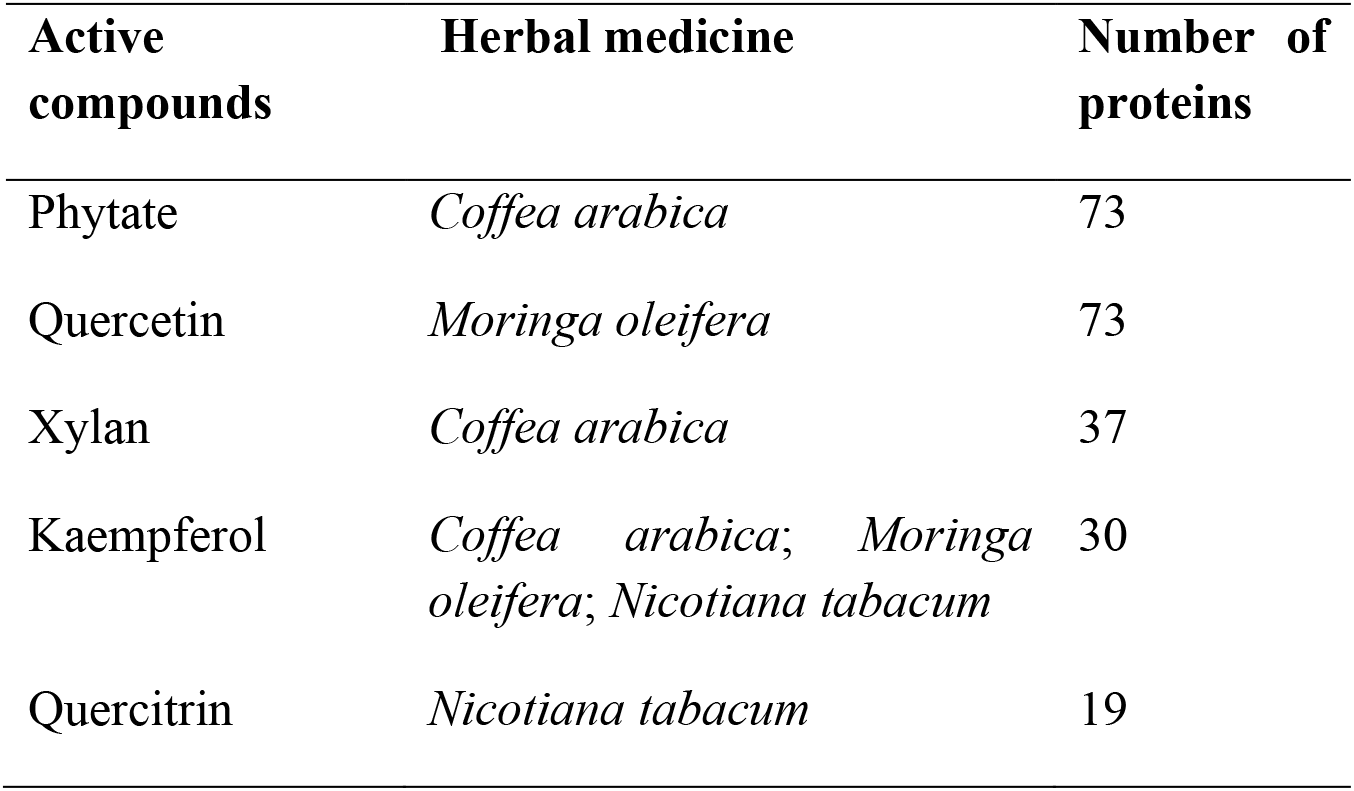
Top five active compounds with a high number of the protein target

Moreover, the result showed that CRABP2, AKR1B1, RORA, TTR, and ACHE were the most protein targeted by various balur active compounds (Table 2). A visualization network of active compounds-protein targets by Cytoscape with 332 nodes and 510 edges as shown in Figure 2. Cellular Retinoic Acid Binding Protein 2 (CRABP2), transporting retinoic acid to the receptor, belongs to the intracellular lipid-binding proteins family and involves in various cancer development. High expression CRABP2 is associated with invasion of lung cancer and malignant peripheral nerve sheath tumors (Fischer-Huchzermeyer et al., 2017; Wu et al., 2019). Aldose reductase (AKR1B1) is one of the members of the aldo-keto reductase (AKR) family taking apart as an antioxidant activity to reduce the production of 4-hydroxy *trans*-2-nonenal (HNE) and glutathionyl-1,4-dihydroxynonene (GS-DHN) leading to macrophage inflammation (Frohnert et al., 2014). Retinoid-related orphan receptor alpha (RORA) is a nuclear receptor protein family that has a gene expression regulation role. A study described that it has potential as a treatment for chronic inflammatory diseases by upregulating IκBα expression (Delerive et al., 2001). Transthyretin (TTR) is an extracellular protein transporting thyroid hormone and retinol-binding. High expression of TTR induces reactive nitrogen species, this can be a biomarker for oxidative stress related to neurodegenerative diseases (Fong & Viera, 2013; Sharma et al., 2009). Acetylcholinesterase (ACHE), an enzyme found in muscles and nerves postsynaptic neuromuscular junctions, is reported that downregulation is responsible for oxidative stress (Liu et al., 2017; Trang & Khandhar, 2020).

**Table 2.** Top five targeted protein by active compounds

| Protein target | Active compounds | Number of active compounds | Herbal medicine |
| --- | --- | --- | --- |
| CRABP2 | Alpha-tocopherol; Campestanol; Caprinic-acid; Cholesterol; Cycloeucalenol; Gamma-sitosterol; Linolenic-acid; Beta-carotene; Cycloartenol; Ergosterol; Flavoxanthin; Lutein; Neoxanthin; Violaxanthin | 14 | *Coffea arabica*; *Nicotiana tabacum* |
| AKR1B1 | 3,4-dicaffeoyl-quinic-acid; 3,5-dicaffeoyl-quinic-acid; 4,5-dicaffeoyl-quinic-acid; Caffetannic-acid; Chlorogenic-acid; Dihydrositosterol; Isochlorogenic-acid; Phytate; Astragalin; Isoquercitrin; Kaempferol; Quercitrin; Rutin; Quercetin | 14 | *Coffea arabica*; *Moringa oleifera*; *Nicotiana tabacum* |
| RORA | Campestanol; Campesterol; Cholestanol; Cholesterol; Citrostadienol; Cycloeucalenol; Lanosterol; Phytosterols; Stigmasterol; Cycloartenol; Ergosterol | 12 | *Coffea arabica*; *Nicotiana tabacum* |
| TTR | Alpha-tocopherol; Campestanol; Cycloeucalenol; Gamma-tocopherol; Squalene; Beta-carotene; Cycloartenol; Ergosterol; Neophytadiene; Neoxanthin; Violaxanthin | 11 | *Coffea arabica*; *Nicotiana tabacum* |
| ACHE | Caffeine; M-cresol; P-cresol; Astragalin; Isoquercitrin; Kaempferol; Nicotiflorin; Quercitrin; Rutin; Quercetin | 10 | *Coffea arabica*; *Moringa oleifera*; *Nicotiana tabacum* |

**Figure 2.**
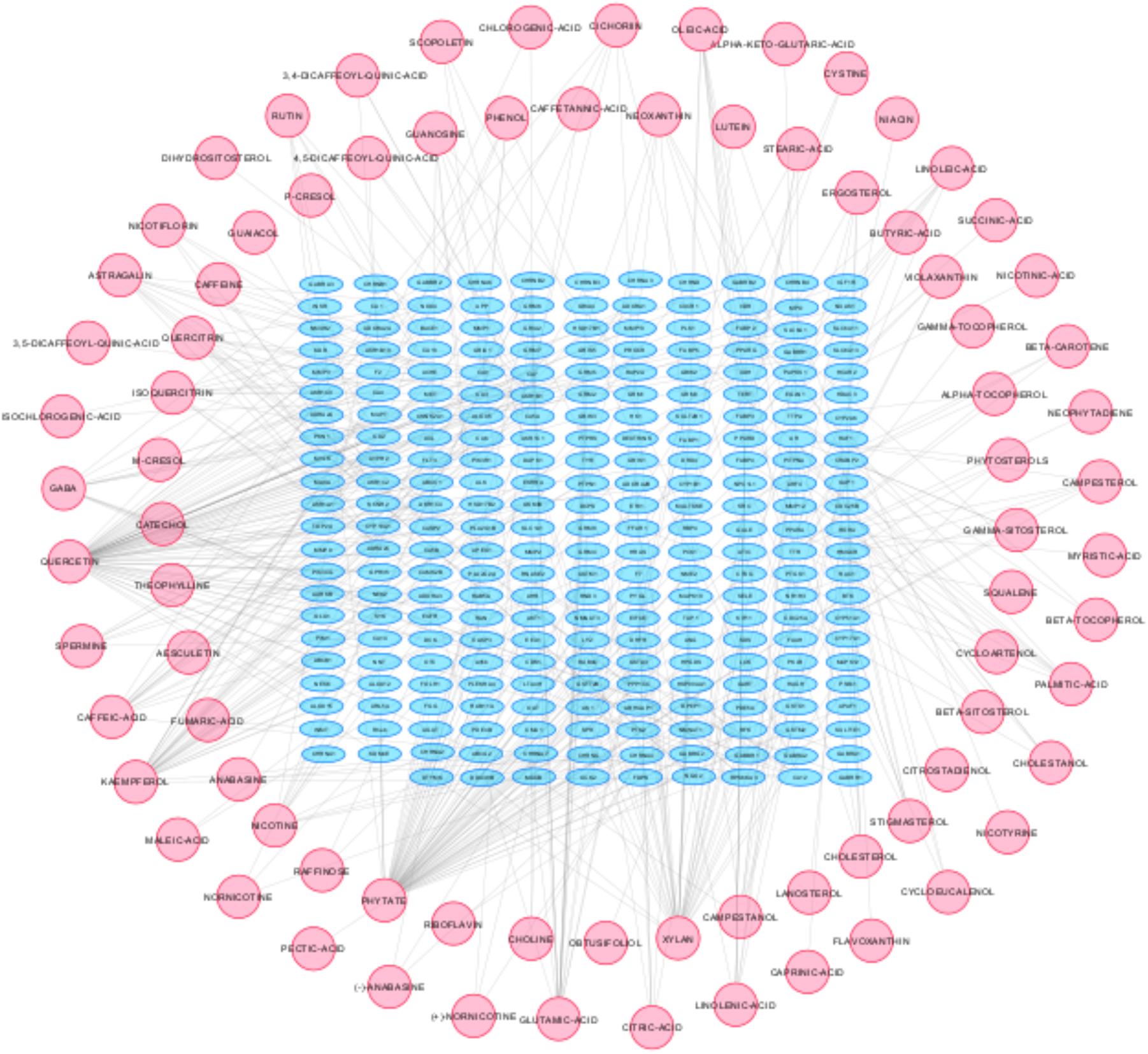
The network of active compounds-target proteins

The result of GO described that from all of the targeted proteins there were 7 molecular functions, 17 biological processes, and 82 pathways which the highest score respectively was catalytic activity, cellular process, and gonadotropin-releasing hormone receptor pathway, Figure 3. Gonadotropin-releasing hormone (GnRH) conducts to lead activation of lipid-derived messengers and mitogen-activated protein kinases, control genes transcription process, and regulate gonadotropin secretion (Bjelobaba et al., 2018). GnRH can upregulate NOS and SOD, enzymes for producing free radicals (Barabutis & Schally, 2008; Perrett & McArdle, 2013). It has been reported that the activation of GnRH for releasing luteinizing hormone (LH) is regulated by nitric oxide (NO), LH will be released when the number of NO is increased (Perrett & McArdle, 2013). However, on another side, multiple studies approved that GnRH correlated with various cancers development (Gründker & Emons, 2017).

**Figure. 3.**
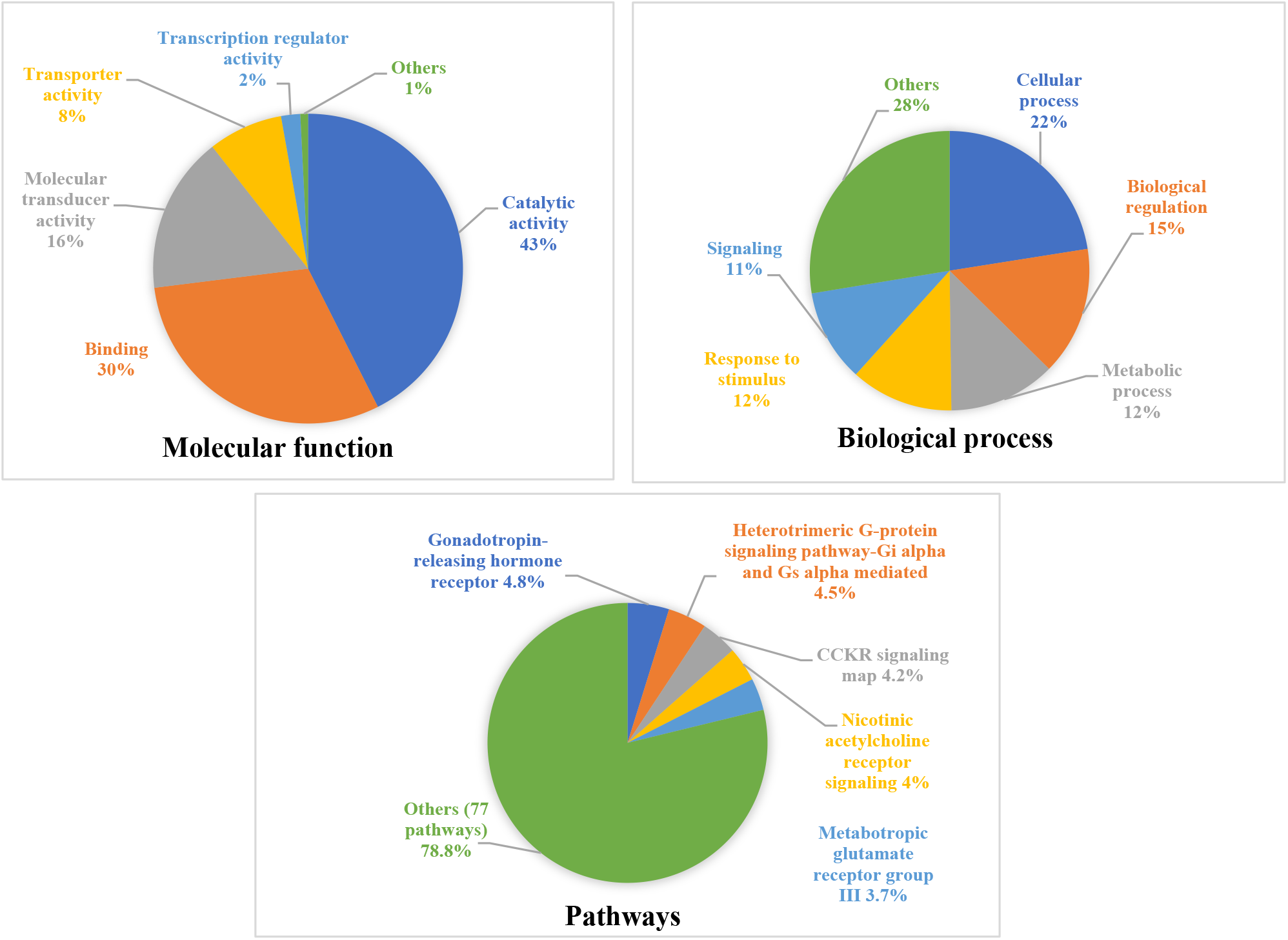
The result of Gene Ontology analysis

In target protein-predictive diseases, 29 predictive diseases were found based on the KEGG PATHWAY database. Cancer was the most disease, then it was followed by, nervous system disease, inherited metabolic disease, congenital malformation, and hematologic disease (Table 3). A network model of target proteins-predictive diseases by Cytoscape generating 125 nodes and 144 edges as shown in Figure 4. According to this result, cancer is a major disease targeted by *Coffea arabica, Moringa oleifera,* and *Nicotiana tabacum* active compounds in balur therapy. One of the cancer risk factors is free radical triggering DNA mutation and damage (Dreher & Junod et al., 1996). However, medical treatment by chemotherapy and radiation can induce to enhance the free radicals in the patient’s body, thus it will be more severe and recurrence (Singh et al., 2018; Mahvi et al., 2018).

**Table 3.** Targeted proteins-predicted disease

| Predicted Disease | Number of target proteins-disease | Predicted Disease | Number of target proteins-disease |
| --- | --- | --- | --- |
| Cancer | 37 | Ribosomopathy | 2 |
| Nervous system disease | 30 | Lung disease | 2 |
| Inherited metabolic disease | 21 | Chromosomal abnormality | 2 |
| Congenital malformation | 20 | Immune system disease | 2 |
| Hematologic disease | 17 | Infectious disease | 1 |
| Mental and behavioral disorder | 13 | Skin disease | 1 |
| Endocrine diseases | 13 | Digestive system disease | 1 |
| Metabolic diseases | 9 | Skeletal diseases | 1 |
| Neurodegenerative disease | 9 | Kidney disease | 1 |
| Musculoskeletal disease | 6 | Liver disease | 1 |
| Cardiovascular disease | 4 | Skin and connective tissue disease | 1 |
| Urinary system disease | 4 | Eye disease | 1 |
| Primary immunodeficiency | 4 | Mental retardation | 1 |
| Vascular disease | 3 | Liver disease | 1 |
| Reproductive system disease | 2 |  |  |

**Figure 4.**
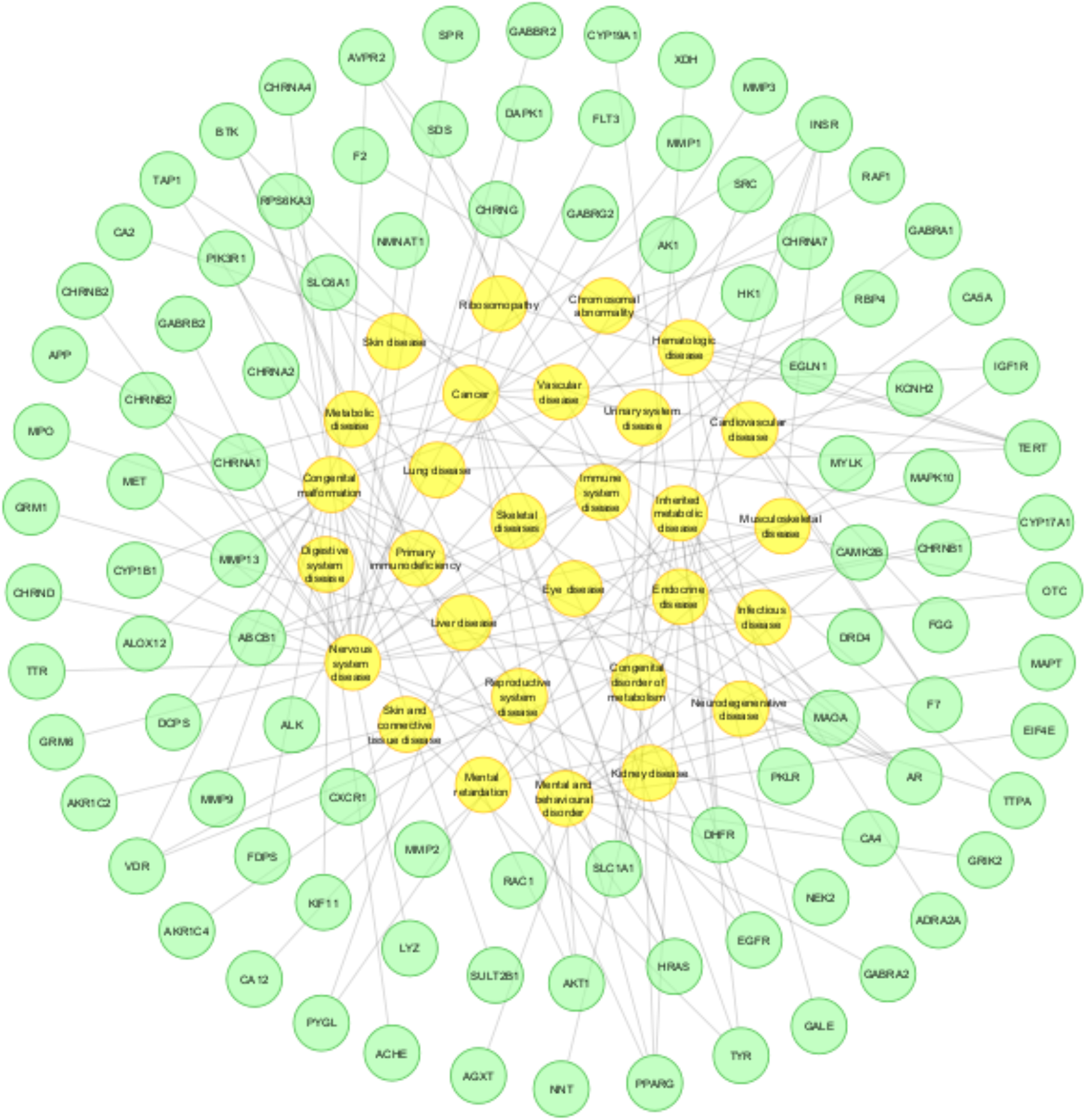
A network of target proteins-predictive disease; green circle: target proteins, yellow circle: predictive disease

Based on these results, it can be concluded that major active compounds of *Coffea arabica, Moringa oleifera* and *Nicotiana tabacum* used in balur played as multiple targeted proteins compounds to reduce excess free radicals in the human body, thus it can use for multiple diseases causing free radicals. However, this study was only from a bioinformatics perspective and it should be to explore other science viewpoints for understanding completely the mechanism of balur. Furthermore, we are in the Research Institute of Free Radicals trying to examine balur in multiple basic science viewpoint to acknowledge balur in global and detail.

## Notes

### Competing Interest Statement

The authors have declared no competing interest.

